# First identification of trombiculid mites (Acari: Trombiculidae) on rodents captured on Chiloé Island, an endemic region of scrub typhus in southern Chile

**DOI:** 10.1101/699512

**Authors:** Gerardo Acosta-Jamett, Esperanza Beltrami, María Carolina Silva de La Fuente, Constanza Martinez-Valdebenito, Thomas Weitzel, Katia Abarca

## Abstract

**Background:** Scrub typhus in an emerging vector-borne zoonosis, caused by *Orientia* spp. and transmitted by larvae of trombiculid mites, called chiggers. It mainly occurs within a certain region of the Asia-Pacific, called tsutsugamushi triangle, where rodents are known as the most relevant hosts for the trombiculid vector. The disease has recently been discovered on Chiloé Island in southern Chile. Still, the reservoir(s) and vector(s) of the scrub typhus outside Asia-Pacific are unknown. The aim of the present work was to study the prevalence of chiggers on different rodent species captured in sites identified as probably hot spots of scrub typhus on Chiloé Island in southern Chile.

**Methodology/Principal Findings:** During austral summer 2018, rodents were live-trapped in six sites and examined for chigger infestation. During a total of 4,713 trap-nights, 244 rodents of seven species were captured: the most abundant was *Abrothrix olivacea*. All study sites were rural areas on Chiloé Island, previously identified as localities of probable human infection with scrub typhus. Chiggers were detected on all seven rodent species with a 55% prevalence rate. Chiggers showed low host specificity and varied according to site specific host abundance. We identified trombiculids of three genera. *Colicus* was the most abundant chigger (93%), prevalent in five of six sites, followed by *Quadraseta* (7%) and *Paratrombicula* (7%), which were in only one site. Infestation rates showed site specific differences, which were statistically different using a GLM model with binomial errors.

**Conclusions/Significance:** This study firstly reports the presence of different rodent-associated chigger mites in a region with endemic scrub typhus in southern Chile. *Colicus* and two other genera of mites were found with high infestation rates in sites previously identified as hot spots of scrub typhus, suggesting their role as vectors and reservoirs of this emerging zoonosis in South America.

**Author Summary:** Scrub typhus is a chigger-transmitted zoonotic infection, which is endemic in the tsutsugamushi triangle in Asia-Pacific. Recently, a first focus of scrub typhus in South America has been confirmed on Chiloé Island in southern Chile. Still, the vectors of scrub typhus in this region remain unknown. We undertook a survey to study the prevalence of chiggers on different rodent species in areas identified as probable hot spots of scrub typhus on Chiloé Island. The study showed that 55% of rodents were infested by trombiculids. Three chigger genera were identified, of which *Colicus* was the most abundant. Chiggers showed low host specificity, but spatial differences. This first demonstration of rodent-associated chigger mites in hot spots of scrub typhus suggests their possible role as vectors of this infection in Chile.

## Introduction

Scrub typhus is a zoonotic disease caused by bacteria of the genus *Orientia*, which causes significant morbidity and mortality [1]. The disease was previously thought to be restricted to a certain region, known as the tsutsugamushi triangle, in Asia-Pacific, but recent cases from the Arabian Peninsula and southern Chile have called this paradigm into question [2-4]. This is supported by serological data and studies in animals mainly from Africa [5].

In Asia-Pacific, the infection is transmitted by larvae of trombiculid mites commonly called chiggers, which are also the reservoir of the bacteria through transovarial and transstadial transmission. Small vertebrates, usually rodents, serve as main hosts of the chiggers and are a critical part of the epidemiology of scrub typhus in Asia–Pacific [6]. Cases mainly occur in rural areas, where populations of infected trombiculid mites are present with a patchy distribution (mite islands) [1]. Surveillance of chiggers has been used as a proxy for the spatial risk of scrub typhus in humans [7,8]. In Asia-Pacific, bacteria is transmitted by different species of *Leptotrombidium* [9], the transmission by other genera in Korea (*Euschoengastia, Neotrombicula*), Japan (*Schoengastia*), and India (*Schoengastiella*) has been suggested, but remains controversial [10,11]. A key factor to understand the distribution and emergence of scrub typhus in these regions is the knowledge of the local chigger fauna, their rodent hosts, and their interaction with environmental and climatic factors [eg. 12,13].

Recently, the first endemic focus of scrub typhus in South America has been confirmed on Chiloé Island in southern Chile [2,3]. Still, the vectors and reservoirs of scrub typhus in regions outside the tsutsugamushi triangle remain unknown. In Chile, 18 species of trombiculid mites have been reported so far, mostly from reptiles [14-16]. Still, none of them belong to the genus *Leptotrombidium* or other genera associated with scrub typhus, and up to now, no studies regarding the rodent-associated chigger fauna have been performed in scrub typhus endemic regions in southern Chile. The aim of the present work was to study the prevalence of chiggers and analyse their infestation pattern on different rodent species captured in sites, which were identified in previous studies as probable hot spots of scrub typhus on Chiloé Island.

## Methods

### Study sites

Chiloé Island belongs to Los Lagos region in southern Chile and is the second largest island in Chile with an area of 8,394 km2. The climate is oceanic temperate, with mean annual precipitations of 2,090 mm and an average annual temperature of 12°C [17]. The original vegetation is Valdivian temperate rain forest, which has been highly fragmented by clearing for livestock rising and timber extraction [18], and the current matrix involves mainly pastures and secondary scrublands [19].

During the austral summer months of January and February 2018, small mammals were live-trapped at six sites in the northern part of Chiloé Island (Figure 1). Study sites had been identified as possible areas of exposure of scrub typhus cases [3]. All sites consisted of partially cleared forest due to timber activities, with remaining native lower vegetation. Localities were geo-referenced by GPS and located into a digitalized map using QGIS 3.6.

**Figure 1.**
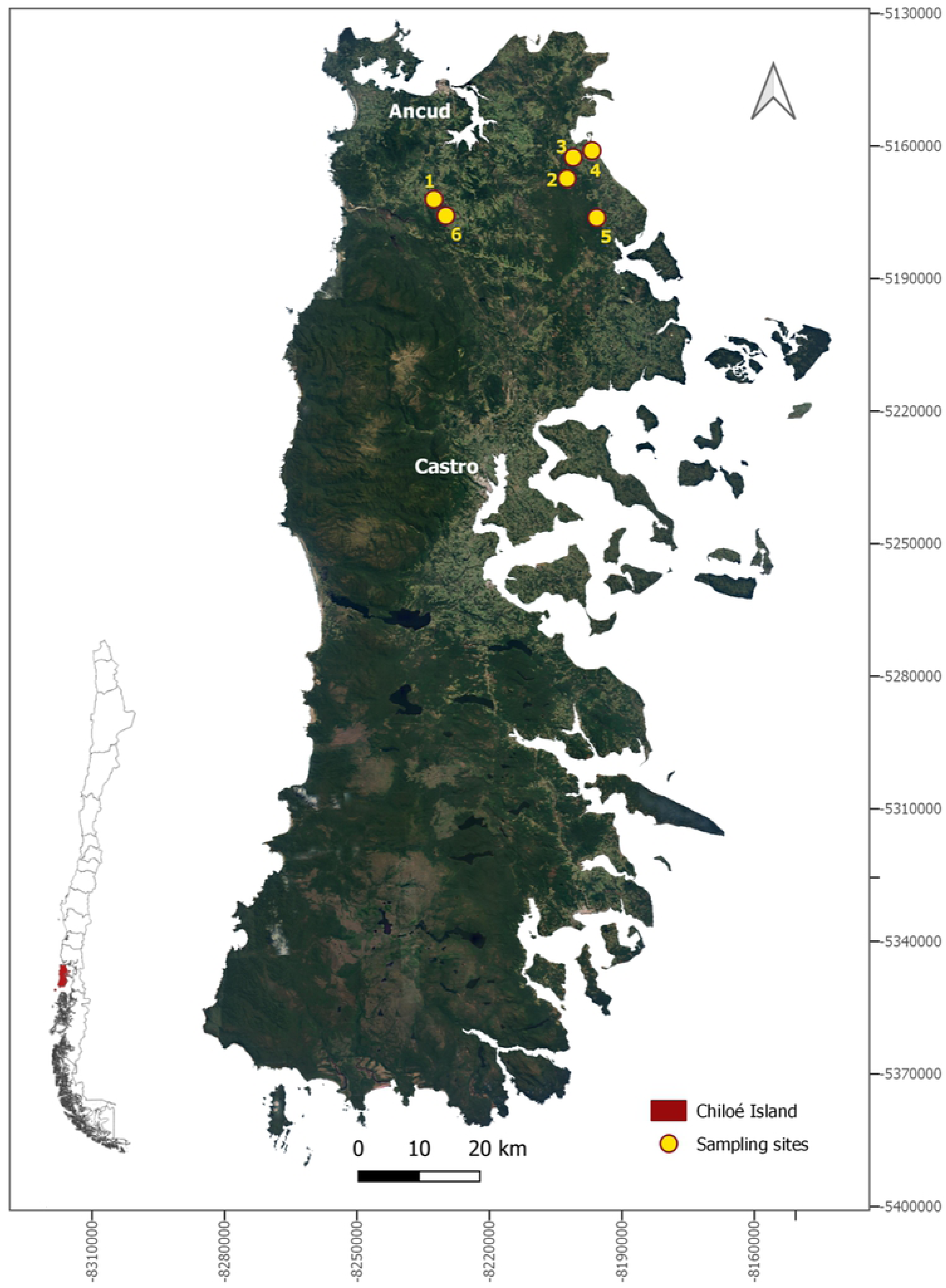
Study area in rural localities of the north-eastern area of Chiloé Island, Los Lagos Region, Chile (Map made in QGIS Geographic Information System. Open Source Geospatial Foundation Project. http://qgis.osgeo.org. Shapes downloaded from an open source from the Biblioteca del Congreso Nacional, Available at https://www.bcn.cl/siit/mapas_vectoriales/index_html)

### Trapping and parasite sampling

A total of 148 to 175 Sherman-like traps (300 × 100 × 110 mm) were set up during four to five consecutive nights at each site. Trapping was operated for a total of 4,713 trap-nights and ranged from 668 to 895 trap-nights per site (average 785.5). Traps were situated ≥5 meters apart and placed under scrub, fallen logs, understory or burrows, baited with oat flakes and vanilla essence, and conditioned both inside and outside with vegetal material to protect animals from cold and rain. Traps were activated at sunset, checked early the next morning, and closed during the day to avoid capturing non-target species. Captured rodents were moved to a central processing tent installed at the sampling site (Figure 2), where they were chemically immobilized using an induction chamber containing cotton embedded with isoflurane (1 ml of isoflurane per 500 ml of chamber volume). After anaesthesia, male and juvenile female rodents were euthanized by cervical dislocation; adult females were marked by haircut and released at the respective capture points. Rodents species were identified by morphological criteria following Iriarte [20]. Each rodent was thoroughly examined for the presence of ectoparasites by brushing the body with a fine comb over a plate covered with water. In adult females with high mite loads, we also conducted a careful scraping of the perianal zone. Chigger mites were collected from the water surface and placed in tubes with 95% ethanol. The skin of euthanized rodents, including ears and perianal zones, was dissected, stored in falcon tubes with 95% ethanol, and later revised for additional chiggers in the entomological laboratory.

**Figure 2.**
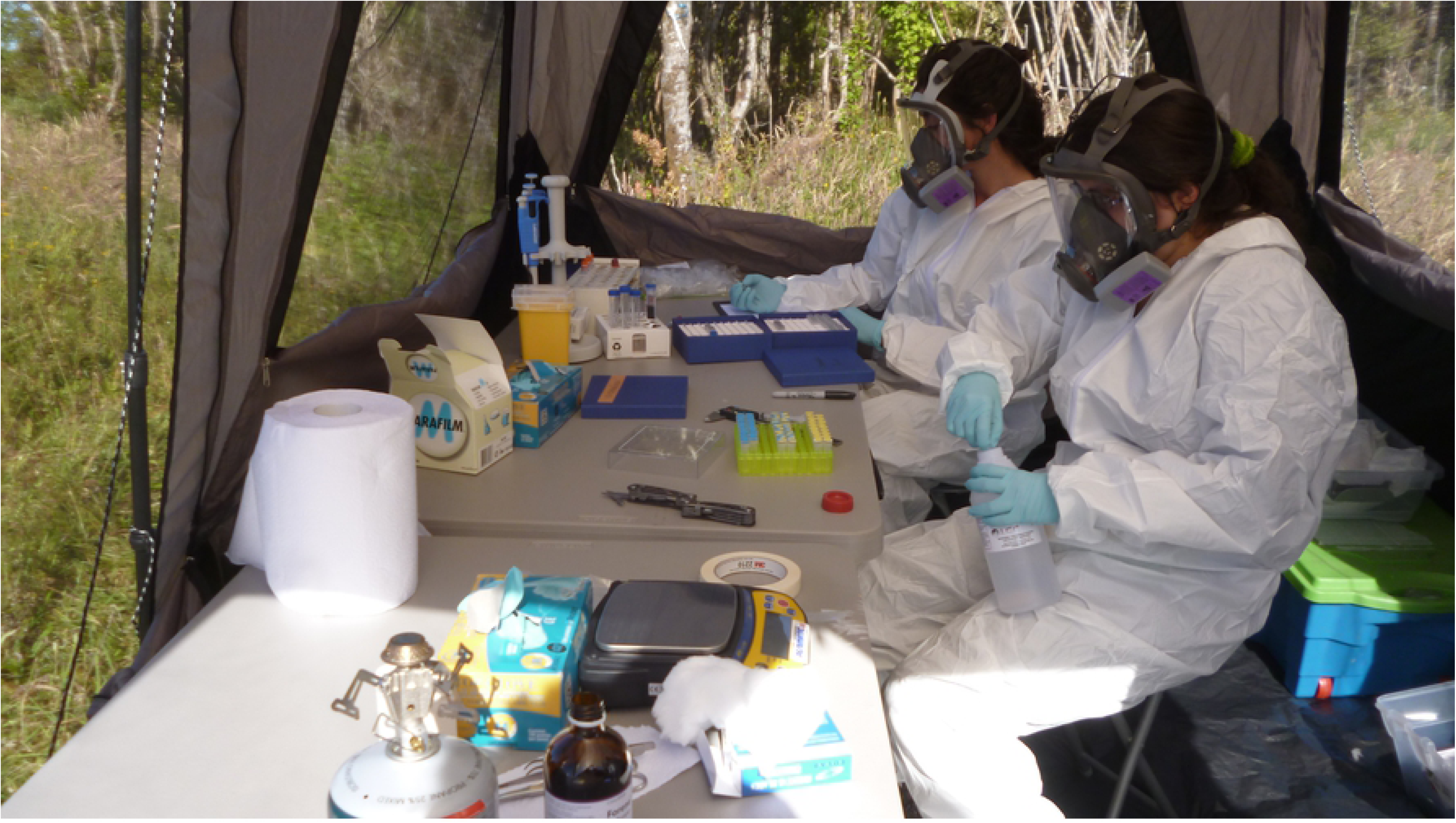
Central processing tent installed at each sampling site. Biosafety conditions during rodent handling were adapted to the risk of Hantavirus cardiopulmonary syndrome.

**Figure 3.**
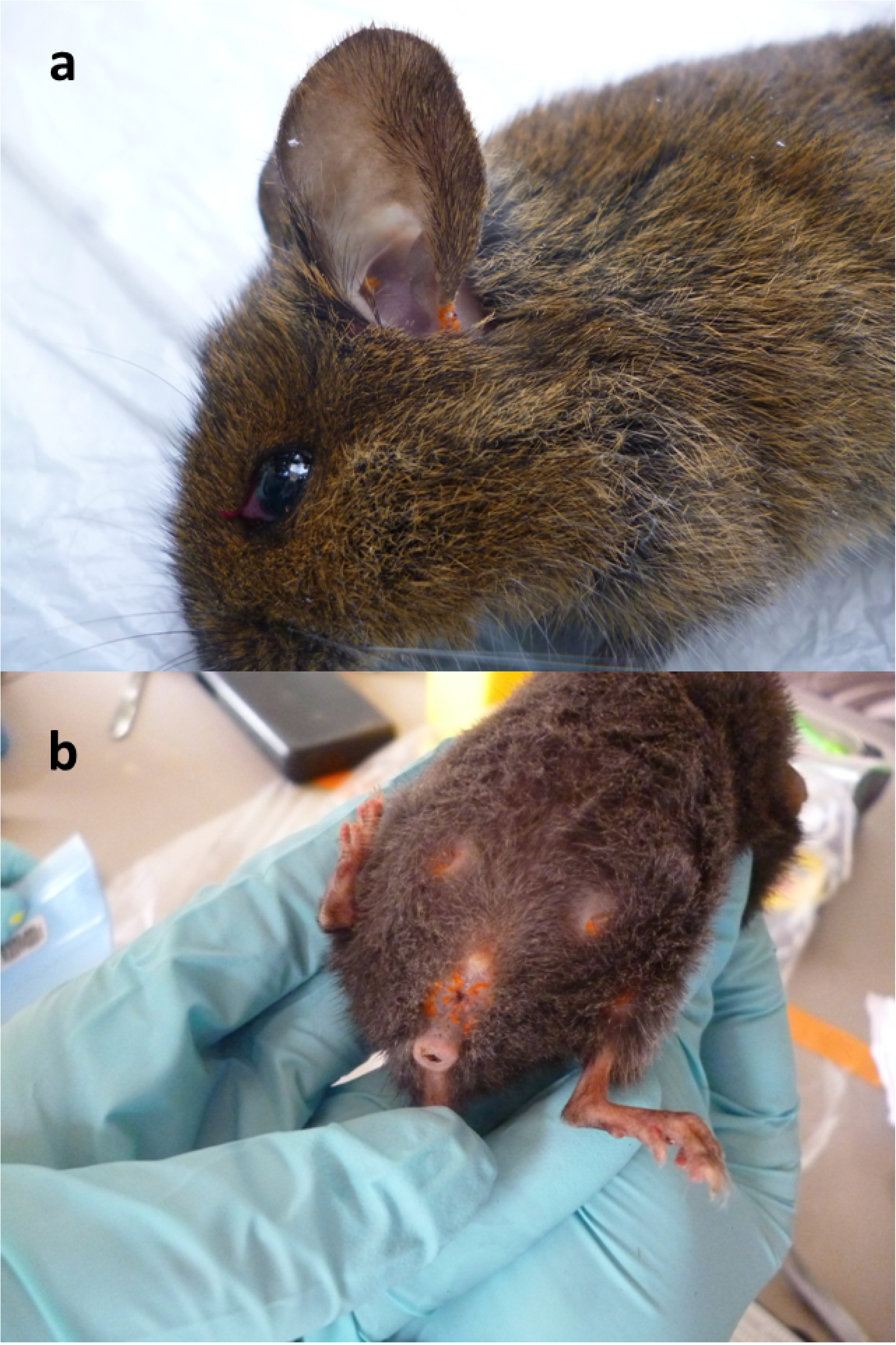
Multiple orange colored trombiculid mites on ear (a) of a *Loxodontomys micropus*, and on genital region and tits of a *Geoxus valdivianus* (b), captured on Chiloé Island.

### Identification of mites

Taxonomic analyses of trombiculid mites were performed at the Laboratorio de Parásitos y Enfermedades de Fauna Silvestre, Universidad de Concepción in Chillán, Chile. Firstly, mite specimens of the individual rodents were pooled by their macroscopic appearance (morphotypes). Preserved samples of skin and ears were checked for additional mites, which were added to the respective pools. One individual of each pool was cleared in Nesbitt’s solution and mounted in Berlese’s medium [21]. Specimens were then identified under an optical microscope (Leica DM 1000 LED) at 400x magnification, following nomenclature and methodology of Brennan & Goff [22].

Southern Chile is endemic for Andes Hantavirus, a rodent-borne virus causing Hantavirus cardiopulmonary syndrome [23]. Therefore, the handling of captured animals strictly followed guidelines of the Centers for Disease Control and Prevention (CDC) [24] and the American Society of Mammalogists [25] for such regions. Personal protective equipment included masks with HEPA filters as well as disposable gowns and gloves (Figure 2).

### Ethics statement

The study also adhered to the guidelines from the American Veterinary Medicine Association [26] and American Society of Mammalogists for the use of wild mammals in research and education [27]. The animal protocol used in this study was approved by the Chilean Animal Health Service (permit number 7034/2017), and by the Scientific Ethics Committee for the Care of Animals and the Environment, Pontificia Universidad Católica de Chile (N°160816007, 07-Nov-2017). All members of the field team were advised and clinically followed for five weeks postexposure for signs and symptoms of scrub typhus and hantavirus; chemoprophylaxis with doxycycline was not used.

### Data analysis

A descriptive analysis of the rodent host community and its infestation pattern with ectoparasites was carried out. The percentage of infected individuals with trombiculid mites per species was estimated. Then, we performed a Generalised Lineal Model (GLM) with binomial errors to assess effects of rodent species, and sites on the chigger prevalence using R, version 3.4.1 [28].

## Results

Of the captured animals, 244 rodents were included, belonging to seven species, most abundantly *Abrothrix olivacea* and *Geoxus valdivianus* (Table 1). Among the trapped animals, 55% (133/244) were infested with trombiculid mites. Other collected ectoparasites included ticks, fleas, and non-trombiculid mites (data not presented). Chigger infestation was observed among all rodent species, with prevalence rates ranging from 77% in *G. valdivianus* to 32% in *Irenomys tarsalis*, without significant species-specific differences (GLM, p>0.05). Overall, the most abundant species (*Abrothrix olivacea*) represented 77% (102/133) of all infected animals.

**Table 1.**
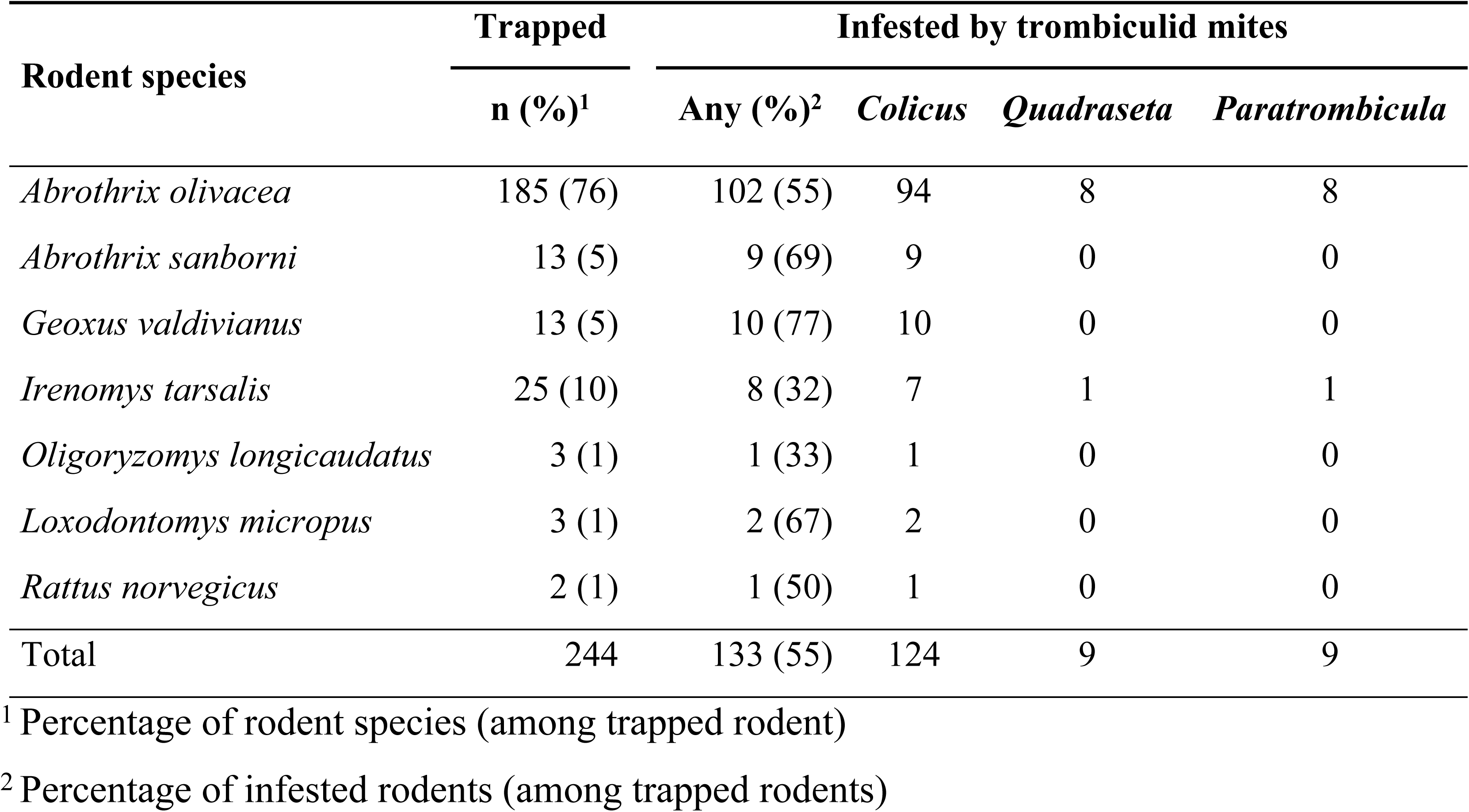
Infestation with different genera of trombiculid mites in 244 rodents captured between January and February 2018 on Chiloé Island.

Among the collected trombiculids, three morphotypes were observed, which were identified as *Colicus* sp., *Quadraseta* sp., and *Paratrombicula* sp.; details of the detected species will be published elsewhere. *Colicus* sp. was predominating (93% of infested rodents) and occurred on all rodent species and in all sites except Site 3, whereas the other two species co-parasitized on few individuals only in Site 3 (Table 2). The overall prevalence of chigger mites in different sites showed significant variations and ranged from 25% to 78% (Table 2). These spatial differences were also present in a Generalized Lineal Model, demonstration that it was less likely to find positive rodents in Sites 3 and 5 than in other sites (Table 3).

**Table 2.**
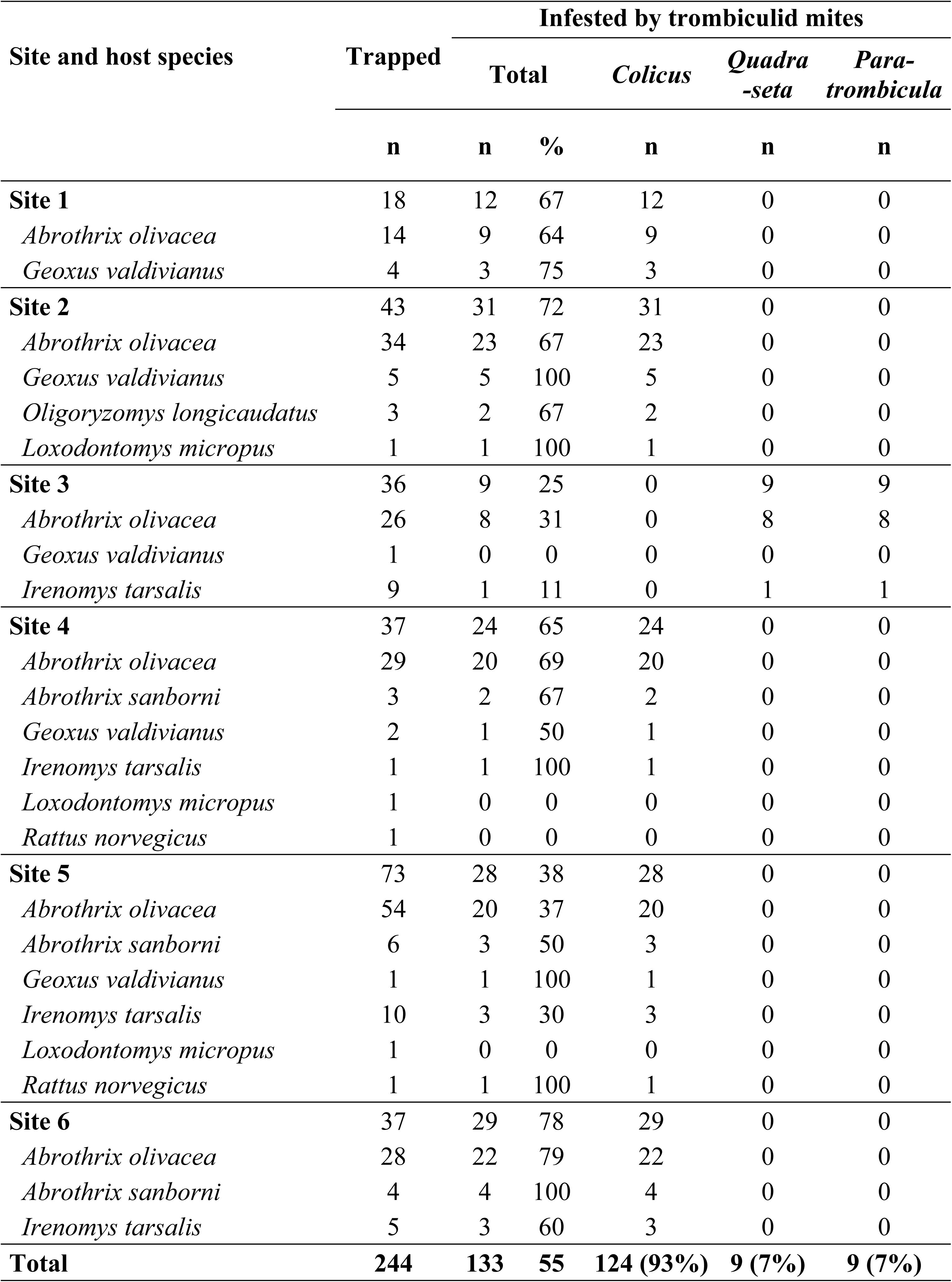
Prevalence of infestation with trombiculid mites per site and rodent species, captured between January and February 2018 on Chiloé Island.

**Table 3.** Generalized Lineal Model with binomial error indicating trapping site as a factor for trombiculid infestation in rodents (n=244) on Chiloé Island.

| Sites | OR* | IC95% | p |
| --- | --- | --- | --- |
| 1 | 1.00 |  |  |
| 2 | 1.29 | 0.39 - 4.22 | 0.672 |
| 3 | 0.17 | 0.05 - 0.57 | 0.005 |
| 4 | 0.92 | 0.28 - 3.03 | 0.895 |
| 5 | 0.31 | 0.10 - 0.92 | 0.035 |
| 6 | 1.81 | 0.52 - 6.35 | 0.353 |

\*OR=Odd ratio

## Discussion

The presented study aimed to explore the trombiculid fauna on Chiloé Island, an endemic area of scrub typhus in southern Chile. Although scrub typhus in Asia-Pacific is transmitted by chigger mites, in Chile and other possible endemic regions, the vector is unknown. Interestingly, a study from 2018 found first molecular evidence for *Orientia chuto* infection in chiggers collected from a rodent in Kenya [29], another region outside the tsutsugamushi triangle, where scrub typhus might occur as suggested by two recent retrospective surveys, detecting antibodies to *Orientia* spp. in 3% to 5% of febrile patients [5,29]. In the endemic regions in Asia-Pacific, the disease is transmitted by different *Leptotrombidium* species; however, mites of this genus are not endemic in Chile or any other region of the Neotropics [30]. Since the first Chilean scrub typhus patient suffered several terrestrial leech bites prior to his infection, it was speculated that *Orientia* was leech-transmitted [2]. This hypothesis cannot be corroborated by our data. None of the cases diagnosed by our group reported leech bites, at least one had symptoms compatible with trombiculidiasis, and most had nature activities with increased risk of arthropod exposure [3,31,32], suggesting that a transmission by chiggers is more likely.

The study was conducted during the typical scrub typhus season in southern Chile, i.e. the summer months of January and February, and locations were chosen according to our previous studies as possible hot spots of *Orientia* transmission [3,33]. We could demonstrate that rodent-associated chigger mites were present in all those sites. The three detected trombiculid genera have been described before in the Neotropics. The genus *Colicus* was most abundant and parasitized all of the captured rodent species. It currently includes 18 species found in rodents, bats, marsupials, and carnivores, from Panama to Argentina [34-40]. *Quadraseta* and *Paratrombicula* were less prevalent. *Quadraseta*, which comprises 14 species found on rodents and birds [36,37,39-44], has not been reported previously in Chile. *Paratrombicula* includes six species isolated from lizards and rodents; two of those (*P. chilensis* and *P. goffi*) have been described in central Chile, both on lizards [14,15,45]. Our preliminary analyses suggest that all three trombiculids found in this study represent new species; further confirmation and detailed morphological description will be presented elsewhere.

Rodents are a main reservoir of chiggers and important determinant of the distribution of scrub typhus in endemic regions in Asia [6]. In our study, *A. olivacea* was the most abundant rodent species and also represented the highest (absolute) number of infested rodents. This species is a typical inhabitant of Valdivian temperate forests; showing a large numerical response during bamboo blossom (*Chusquea* spp.) [46-48]. Other less abundant host species were infested in similar rates, indicating low host-specificity of trombiculids. This is in accordance with studies from Asia-Pacific, where feeding on small mammals was non-specific and depended on host species abundance in the community [1,7,8].

Although chigger mites were detected on rodents in all six surveyed areas, the prevalence rates differed geographically. Variations in the distribution of trombiculids are a known phenomenon; in fact, within suitable habitats, the mites usually have a patchy distribution, forming so called “mite islands” [9,49]. Our findings might indicate a high prevalence of chiggers on Chiloé Island, although the selection of sites as probable “hot spots” of exposure to *Orientia* spp. could be a bias towards overestimation. The infestation rates reported in this study are comparable to those reported in the Asia-Pacific region where prevalence rates on small mammals ranged from 45% to 95% [7,8,50,51].

Trombiculid mites live in moist soil covered with vegetation and are mostly found in grassy and weedy areas [eg. 49]. Optimal living conditions depend on various factors such as air humidity, soil composition, temperature, and light intensity. Habitat fragmentation seems to affect mite survival by modifying their ecological niche [52]. Recently, a large-scale research found higher infestation (prevalence, mean abundance, and intensity) with vector mites on small mammals in areas with lower biodiversity compared to those with higher biodiversity [53]. As documented in various studies, forest fragmentation on Chiloé Island reduces the biodiversity of small mammals, non-raptorial birds, and the predator assemblage and increases the abundance of generalist species [54-57]. In a similar manner, fragmentation might affect rodent populations and their associated trombiculid ectoparasites.

This first demonstration of rodent-associated chigger mites in probable hot spots of scrub typhus suggests that chiggers might serve as vectors of this infection in Chile. The study detected high chigger prevalence in the summer season, during which up to now all cases of scrub typhus in Chiloé have occurred [58]. However, final prove of the vector competence of the detected trombiculid mites in this new endemic region requires further studies. Molecular analyses to detect *Orientia* DNA within the collected chigger and rodent specimens are currently under way. To understand the risk of human exposure to trombiculid mites, further investigations are necessary, which should include environmental, anthropogenic, and climatic variables influencing the epidemiology of these potential vectors in southern Chile.

## Conclusions

Our study firstly documented the prevalence of rodent-associated trombiculid mites on Chiloé Island, a region endemic for scrub typhus in South America. Three different mite genera were identified; the neotropical genus *Colicus* was the most abundant. Overall, we detected a high rate of chigger infestation independent of host species, but with significant spatial variations.

## Acknowledgements

We acknowledge the Comisión Nacional de Investigación Científica y Tecnológica (CONICYT) for the PhD scholarships of Maria Carolina Silva-de la Fuente and Esperanza Beltrami. We also thank Maira Riquelme and Gunther Heyl for their assistance during field work.

## Author’s Contributions

GAJ, TW and KA designed the study and GAJ and EB wrote the manuscript. EB and GAJ collected the data. MCSF carried out the chigger identification. TW, KA and CMV reviewed and edited the manuscript.

## Data Availability

The authors confirm that all data underlying the findings are fully available without restriction. All relevant data are within the paper and its Supporting Information files.

